# Hydrogel mechanics regulate fibroblast DNA methylation and chromatin condensation

**DOI:** 10.1101/2022.12.13.520341

**Authors:** Jenna L. Sumey, Peyton C. Johnston, Abigail M. Harrell, Steven R. Caliari

## Abstract

Cellular mechanotransduction plays a central role in fibroblast activation during fibrotic disease progression, leading to increased tissue stiffness and reduced organ function. While the role of epigenetics in disease mechanotransduction has begun to be appreciated, there is little known about how substrate mechanics, particularly the timing of mechanical inputs, regulate epigenetic changes such as DNA methylation and chromatin reorganization during fibroblast activation. In this work, we engineered a hyaluronic acid hydrogel platform with independently tunable stiffness and viscoelasticity to model normal (storage modulus, G’ ∼ 0.5 kPa, loss modulus, G’’ ∼ 0.05 kPa) to increasingly fibrotic (G’ ∼ 3.5 and 8 kPa, G’’ ∼ 0.05 kPa) lung mechanics. Human lung fibroblasts exhibited increased spreading and nuclear localization of myocardin-related transcription factor A (MRTF-A) with increasing substrate stiffness within 1 day, with these trends holding steady for longer cultures. However, fibroblasts displayed time-dependent changes in global DNA methylation and chromatin organization. Fibroblasts initially displayed increased DNA methylation and chromatin decondensation on stiffer hydrogels, but both of these measures decreased with longer culture times. To investigate how culture time affected the responsiveness of fibroblast nuclear remodeling to mechanical signals, we engineered hydrogels amenable to *in situ* secondary crosslinking, enabling a transition from a compliant substrate mimicking normal tissue to a stiffer substrate resembling fibrotic tissue. When stiffening was initiated after only 1 day of culture, fibroblasts rapidly responded and displayed increased DNA methylation and chromatin decondensation, similar to fibroblasts on static stiffer hydrogels. Conversely, when fibroblasts experienced later stiffening at day 7, they showed no changes in DNA methylation and chromatin condensation, suggesting the induction of a persistent fibroblast phenotype. These results highlight the time-dependent nuclear changes associated with fibroblast activation in response to dynamic mechanical perturbations and may provide mechanisms to target for controlling fibroblast activation.

## 1. Introduction

Cellular mechanotransduction, or the process by which cells interpret and translate mechanical cues from their surrounding microenvironment into biochemical signals, has implications in a variety of cellular processes such as homeostasis, differentiation, proliferation, and apoptosis^1-4^. Additionally, mechanical cues from the extracellular matrix (ECM) directly drive aberrant cellular behaviors in tissue stiffening pathologies like fibrosis^5, 6^. Fibrosis is characterized by excessive and irregular ECM deposition, leading to increased tissue stiffness and reduced viscoelasticity, which impairs organ function and ultimately results in death^5, 7-11^. A positive feedback loop between the matrix-depositing fibroblasts and the surrounding ECM drives fibrogenesis, including activation of cytoskeletal signaling pathways for cell spreading and elongation, as well as fibroblast activation into fibrogenic myofibroblasts^4, 12-14^.

Over the past few decades, hydrogel biomaterial platforms enabled researchers to investigate cell-matrix communication in physiologically-relevant settings, unlike traditional tissue culture plastic, without the presence of extraneous and confounding biological signals present *in vivo*. This resulted in the discovery that mechanical cues, like substrate elastic modulus or stiffness, can directly promote fibroblast activation^15, 16^. Studies supporting this notion evaluated cells cultured on increasingly stiff matrices and found greater cell spreading, actin stress fiber formation, and nuclear translocation of transcription factors involved in cellular activation^3, 17-23^. Furthermore, more complex mechanical cues like viscoelasticity, or exhibition of both solid- and liquid-like properties with time-dependent strain response, have also been found to play key regulatory roles in cell behavior. While healthy tissues like lung display low stiffness and higher viscoelasticity, aging and/or fibrotic ECM becomes increasingly stiff and more elastic over time^5, 8, 10, 24,25^. Recent studies found that viscoelastic hydrogels supported reduced cell spreading and nuclear translocation of transcriptional factors as a result of lower cellular contractility compared to elastic hydrogels of similar stiffness^26-33^.

While it’s well understood that mechanical cues drive pro-fibrotic cytoskeletal behaviors including spreading, actin stress fiber organization, and focal adhesion formation, there is comparatively little known about how these mechanical signals are translated into observed phenotypic responses, as well as how the timing of mechanical signals regulates the persistence of fibrotic phenotypes. To address this point, epigenetic mechanisms have recently been investigated for their contributions to cellular mechanotransduction. The nucleus serves a central role in mechanotransduction through its cytoskeletal adhesions; furthermore, the structural organization within the nucleus, such as chromatin organization, provides the cues for transcriptional events during cell state transitions^34-36^. Epigenetic alterations involve changes in chromatin organization, as well as DNA or histone accessibility, which result in gene regulation and subsequent changes in cell state or differentiation^35^. Seminal studies elucidated the role of chromatin remodeling, with initially greater accessibility followed by condensation, in persistently phenotypically-activated cells, and others have shown that this response is a result of prolonged exposure to stiff substrates^37-42^. These observations have implications in fibrotic disorders where persistently activated cells drive fibrogenesis, and this persistent phenotype is supported by underlying nuclear mechanisms. Similarly, DNA methylation has been shown to play a crucial role in a number of cellular functions like differentiation, tumorigenesis, and fibrosis^43^. DNA methylation is mediated through DNA methyltransferases that covalently attach methyl moieties to residual cytosines within cytosine-guanine (CpG) islands. DNA methylation can regulate cellular differentiation and is sensitive to mechanical cues, with increased global methylation observed on stiffer substrates^44, 45^. Therefore, DNA methylation, along with chromatin accessibility, may be crucial regulators of cell fate and responsible for the persistent activation of fibroblasts, although little is known about how DNA methylation changes over time in response to substrate mechanics.

While these studies have highlighted the importance of epigenetics on cell fate and differentiation, less is known about how substrate mechanics regulate the time-dependent effects of DNA methylation in conjunction with chromatin remodeling. We created a pathophysiologically-relevant hydrogel model of normal and increasingly fibrotic lung tissue through combined control of stiffness and viscoelastic mechanical cues to investigate these key nuclear markers, as well as time-dependent effects including increased culture time and response to dynamic substrate stiffening, to explore their implications in the persistence of fibroblast activation and thus, fibrogenesis.

## 2. Materials and Methods

### 2.1 NorHA synthesis

Hyaluronic acid (HA) was modified with norbornene moieties as previously reported^27, 46^. Sodium hyaluronate (Lifecore, 82 kDa) underwent proton exchange using Dowex 50W resin to transform into hyaluronic acid tert-butyl ammonium salt (HA-TBA). The reaction mixture was filtered, titrated to pH 7.05, frozen, then lyophilized to dryness. HA-TBA was then reacted with 5-norbornene-2-methylamine and benzotriazole-1-yloxytris-(dimethylamino)phosphonium hexafluorophosphate (BOP) in dimethylsulfoxide (DMSO) for 2 h at room temperature, then quenched with cold water. The reaction mixture was dialyzed (molecular weight cut off: 6-8 kDa) for 5 days, filtered, then dialyzed for another 5 days prior to freezing and lyophilization. The degree of modification was 30% as determined by ^1^H NMR (500 MHz Varian Inova 500, **Fig. S1**).

### 2.2 β-CD-HDA synthesis

β-cyclodextrin hexamethylene diamine (β-CD-HDA) was synthesized using a previously described method^47^. A solution of p-Toluenesulfonyl chloride (TosCl) dissolved in acetonitrile was added dropwise to aqueous β-cyclodextrin (CD) (5:4 molar ratio of TosCl:CD) at 25°C and allowed to react for 2 h. The reaction mixture was then placed on ice and an aqueous NaOH solution was added dropwise (3.1:1 molar ratio of NaOH:CD) and allowed to react for 30 min at 25°C. Ammonium chloride was added to adjust the pH to 8.5, then the solution was cooled on ice, precipitated with cold deionized (DI) water followed by acetone, and then dried overnight. The CD-Tos product was added to hexamethylene diamine (HDA) (4 g HDA/1 g CD-Tos) in dimethylformamide (DMF) (5 mL DMF/1 g CD-Tos) and allowed to react at 80°C for 12 h under nitrogen. The reaction solution was then precipitated in cold acetone (5 × 50 mL acetone/1 g CD-Tos), washed with cold diethyl ether (3 × 100 mL), and dried. The β-CD-HDA product was confirmed using H^1^ NMR (**Fig. S2**).

### 2.3 β-CD-HA synthesis

β-cyclodextrin-functionalized hyaluronic acid (CD-HA) was synthesized through the reaction of β-CD-HDA to HA-TBA with BOP in anhydrous DMSO^47^. The amidation was performed at 25°C for 2-3 h then quenched with cold DI water and dialyzed against water for 5 d, filtered, and dialyzed for another 5 d. The product was then frozen and lyophilized. The degree of modification was determined as 31% using H^1^ NMR (**Fig. S3**).

### 2.4 Thiolated adamantane peptide synthesis

Solid phase peptide synthesis was carried out on a Liberty Blue (CEM) peptide synthesizer. A thiolated adamantane-containing peptide (Ad-KKKCG) was synthesized on Rink Amide MBHA high-loaded (0.78 mmol/g) resin. Following the reaction, the peptide was cleaved in a cocktail containing 95% trifluoracetic acid, 2.5% triisopropylsilane, and 2.5% water for 3 h. The solution was then precipitated in cold ether, dried, resuspended in water, frozen, and lyophilized. Synthesis completion was determined using matrix-assisted laser desorption/ionization (MALDI) mass spectrometry (**Fig. S4**).

### 2.5 Hydrogel formation

Viscoelastic hydrogel films were crosslinked between an untreated and thiolated glass coverslip (50 μL hydrogel volume, 18 × 18 mm). 5 wt% HA precursor solutions were made by first mixing CD-HA (8 wt% stock solution) with Ad-KKKCG (1:1 molar ratio of CD:Ad) to incorporate the guest-host interactions between Ad-CD before mixing in the remaining components: NorHA (8 wt% stock solution), a thiolated RGD peptide (GCGYG<u>RGD</u>SPG, 1 mM, GenScript), lithium acylphosphinate photoinitiator (LAP, 1 mM), and dithiothreitol crosslinker (DTT, 0.06, 0.2, and 0.8 thiol:norbornene ratio for 1.5, 7, and 24 kPa hydrogels, respectively). The solutions were photopolymerized using ultraviolet (UV) light (365 nm, 5 mW/cm^2^) for 2 min. The hydrogels were swelled in phosphate-buffered saline (PBS) overnight at 37°C prior to any experiments.

### 2.6 Secondary hydrogel stiffening

To incorporate secondary network crosslinks, hydrogels initially fabricated with a 0.06 thiol:norbornene ratio were allowed to swell in additional 1 mM LAP and DTT to obtain a total 0.8 thiol:norbornene ratio for 30 min at 37°C. The hydrogels were photocrosslinked under the same conditions as initial fabrication.

### 2.7 Mechanical characterization

Hydrogel mechanics were determined through rheological measurements on an Anton Paar MCR 302 rheometer using a cone-plate geometry (25 mm diameter, 0.5°, 25 μm gap) at 25°C. Hydrogel mechanical properties were tested using an oscillatory time sweep (1 Hz, 1% strain) with a 2 min UV irradiation (365 nm, 5 mW/cm^2^), oscillatory frequency sweep (0.001-10 Hz, 1% strain), and cyclic stress relaxation and recovery (0.1 and 5% alternating strain)^26^.

### 2.8 Cell culture

Human lung fibroblasts (abm hTERT T1015) were used at passage 1 for all experiments. Cells were maintained in media comprised of Dulbecco’s modified Eagle’s medium (DMEM) supplemented with 10 v/v% fetal bovine serum (FBS, Gibco), and 1 v/v% streptomycin/amphotericin B/penicillin at 10,000 μg/mL, 25 μg/mL, and 10,0000 units/mL, respectively (Gibco). Cells were maintained in tissue culture plastic (TCP) flasks to ∼ 80% confluency prior to passaging. For expansion, cells were incubated in 0.025% trypsin-EDTA (Gibco) for 6-8 min. FBS-containing medium was added to inactivate trypsin, and the cell solution was centrifuged at 200 rcf for 5 min. The supernatant was aspirated, and the remaining cell pellet was resuspended with warmed media to a final concentration of 250,000 cells/flask. For *in vitro* experiments, swollen hydrogels were placed in non-tissue culture treated 6-well plates and sterilized under germicidal UV light for 2 h, then incubated in warmed culture media for at least 30 min before seeding cells. Cells were trypsinized from culture plates under the same conditions as TCP expansion and seeded at a density of 2 × 10^3^ cells per hydrogel. For all experiments, culture media was replaced every 2-3 days.

### 2.9 Cell staining, fluorescence imaging, and quantification

For immunocytochemical staining of F-actin, myocardin-related transcription factor A (MRTF-A), and nuclei, cells on hydrogels were rinsed with PBS, fixed in 10% buffered formalin for 15 min, permeabilized in 0.1% Triton X-100 for 10 min, then blocked with 3% bovine serum albumin (BSA) for at least 1 h. Hydrogels were then incubated in primary MRTF-A antibody (mouse monoclonal anti-Mk11 Abcam ab219981, 1:200) at 4°C overnight. The following day, hydrogels were rinsed three times with PBS, incubated in secondary antibodies rhodamine phalloidin (Invitrogen R415, 1:600), to visualize F-actin, and AlexaFluor 488 (goat anti-mouse IgG 1:400) to visualize MRTF-A for 2 h at room temperature. The hydrogels were washed three times with PBS then incubated in DAPI nuclear stain (Invitrogen D1306, 1:10,000) for 1 min. The hydrogels were rinsed two times and stored in the dark at 4°C prior to imaging. To visualize global DNA methylation, cells were fixed and permeabilized as above. Hydrogels were then incubated in 4 N HCl for 8 min, rinsed three times with DI water, and neutralized with 100 mM Tris-HCl for 20 min. Hydrogels were rinsed with DI water, then blocked in 3% BSA for at least 1 h at room temperature. Hydrogels were then labeled with primary 5-methylcytosine (recombinant rabbit monoclonal, ThermoFisher RM231, 1:200) antibody, flipped upside-down, and incubated overnight at 4°C. The following day, the hydrogels were returned upright and rinsed three times in PBS prior to incubating in secondary antibody (AlexaFluor 488 goat anti-rabbit, 1:200) upside down for 2 h at room temperature. Hydrogels were returned upright, rinsed three times with PBS, and incubated with DAPI (1:10,000) for 1 min., rinsed twice more with PBS, then stored at 4°C prior to imaging.

For cell cytoskeletal imaging, images were obtained on a Zeiss AxioObserver 7 inverted microscope at 40x (oil objective, 1.3 numerical aperture). To quantify cell spread area, cell shape index (CSI), and MRTF-A nuclear/cytosolic ratio, a pipeline developed with CellProfiler (Broad Institute, Harvard/MIT) was used. CSI measures the circularity of the cell, where a line and circle are equated to values of 0 and 1, respectively, and was determined using the following formula:

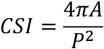

where *A* is the area and *P* is the perimeter of the cell. To image chromatin condensation, cell nuclei were imaged at 63x (oil objective, 1.4 numerical aperture). At least 20 images were taken of each hydrogel (60 total images per experimental group). To quantify the percentage of condensed chromatin (CCP), at least 20 individual nuclei from each hydrogel were obtained using the crop feature within the Zeiss ZEN Imaging software to generate images containing a single nucleus. A gradient-based Sobel edge detection algorithm developed in MATLAB^48^ was used to measure the edge densities within the nuclei to quantify CCP.

### 2.10 Statistical analysis

For mechanical characterization of hydrogels, at least 3 hydrogel replicates were used, and the data are presented as the mean ± standard deviation. One- or two-way analysis of variance (ANOVA) followed by Tukey’s post-hoc tests were completed for all quantitative tests. Cell experiments included at least 3 replicate hydrogels per group. Box plots of single cell data include the total mean/median indicators along with error bars corresponding to the smaller value of either 1.5 X interquartile range or the maximum/minimum value. Data points outside of 1.5 X interquartile range are reported as open circles. Statistical significance is indicated by *, **, ***, or **** pertaining to *P* < 0.05, 0.01, 0.001, or 0.0001, respectively.

## 3. Results

### 3.1 Hyaluronic acid-based hydrogels were fabricated with independently tunable stiffness and viscoelastic mechanical properties

Hydrogels comprised of hyaluronic acid (HA) were formed through the incorporation of covalent and physical crosslinks via thiol-ene click reactions to match mechanical properties of human lung tissue^27^ (**Fig. 1**). Norbornene-modified HA enabled both the formation of a covalently-crosslinked network using dithiol crosslinker (DTT) and physical association of thiolated adamantanes with β-cyclodextrin-functionalized HA via guest-host supramolecular inclusion (**Figs. S1-S4**). Notably, this system enables independent control of hydrogel storage and loss modulus, so that stiffer and more elastic hydrogels could be formed by increasing the number of covalent crosslinks while keeping the physical associations constant (**Fig. 2A**). Rheological analysis performed prior to, during, and following UV light exposure (365 nm, 5 mW/cm^2^) showed the increase in storage and loss moduli as the hydrogels form, then a plateau as the crosslinkers become consumed (**Fig. 2B**). Three stiffnesses with Young’s moduli of *E* ∼ 1.5, 7, and 24 kPa were fabricated to model compliant and increasingly fibrotic lung tissue^49^. Notably, the loss moduli for each hydrogel stiffness group showed no statistically significant differences (**Fig. S5**). Frequency sweeps showed increasing loss moduli for all hydrogel groups, indicating the dissociation of physical guest-host bonds with higher frequencies (**Fig. 2C**). Stress relaxation measurements showed a greater relaxation response for the *E* ∼ 1.5 kPa hydrogels, indicating that these hydrogels are both more viscoelastic (higher G”:G’ ratio) and stress relaxing compared to the two higher stiffness hydrogel groups (**Fig. 2D**).

**Fig. 1:**
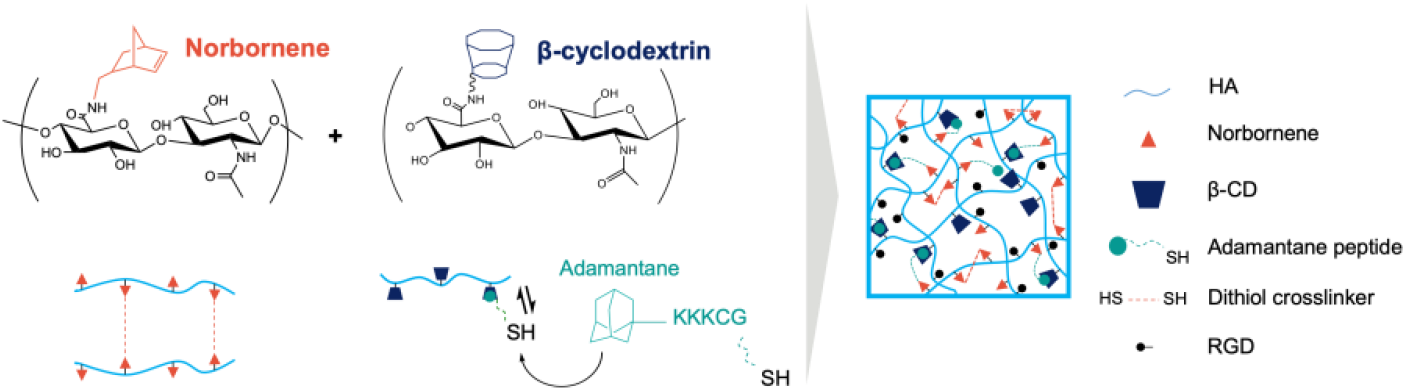
Schematic of hydrogel crosslinking mechanisms. Norbornene-modified hyaluronic acid (HA) and β-cyclodextrin-modified HA were used to form thiolated covalent crosslinks and physical crosslinks with adamantanes, respectively. Thiol-ene click chemistry was used to tether thiolated adamantane moieties in addition to thiolated RGD peptides for cell adhesion.

**Fig. 2:**
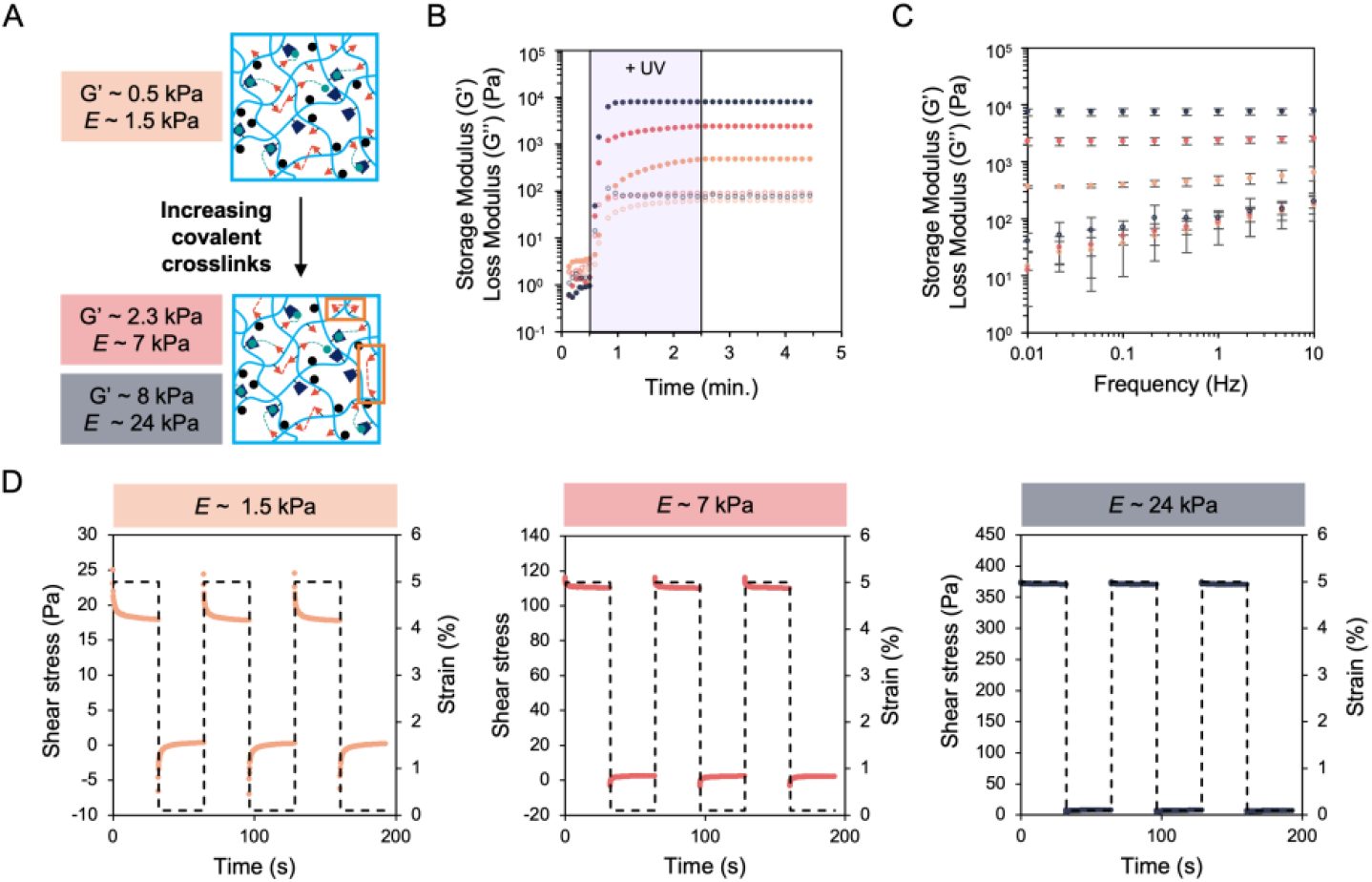
Mechanical characterization of increasingly stiff and elastic hydrogels. A) Schematic of crosslinking chemistry illustrating formation of stiffer hydrogels through the incorporation of increased covalent dithiol crosslinks. B) Rheology showed a loss modulus (G’’, open circles) within an order of magnitude of the storage modulus (G”, closed circles) for the softest, G’ ∼ 0.5 kPa, (orange) viscoelastic hydrogel. Hydrogels with G’ ∼ 2.5 kPa (red) and ∼ 8 kPa (blue) were made with the same loss moduli of the softest hydrogel to form stiffer and more elastic hydrogels mimicking progressively more fibrotic lung tissue. C) Frequency sweeps of the three hydrogel groups illustrated increased loss moduli with increasing frequency. Error bars represent the S.D. of the average of three tests. D) Stress relaxation testing for the three hydrogel groups showed greater relaxation for the softest hydrogel group while the stiffer groups displayed less relaxation. At least three tests were performed per group.

### 3.2 Human lung fibroblasts respond to substrate mechanical cues within 1 day

To probe whether human lung fibroblasts displayed phenotypic sensitivity to hydrogels of increasing stiffness and elasticity, cells were cultured atop the three hydrogel groups, as well as a glass non-hydrogel control group, for a period of 9 days. Within 1 day, fibroblasts exhibited increased formation of F-actin stress fibers and nuclear translocation of MRTF-A with increasing substrate stiffness (**Fig. 3A**). Fibroblasts exhibited greater spreading and elongation (as measured by cell shape index) with increasing stiffness as well (**Fig. 3B, C**). Furthermore, quantification of MRTF-A nuclear localization reported higher values for fibroblasts cultured on stiffer substrates (**Fig. 3D**). Quantification of these same metrics after 9 days showed similar results as day 1, suggesting that fibroblasts rapidly respond to substrate mechanics and do not significantly change their spreading behavior over longer culture lengths (**Fig. S6**). Of note, fibroblast shape and MRTF-A localization does not seem to depend on substrate viscoelasticity at lower stiffnesses, as fibroblasts cultured on *E* ∼ 1.5 kPa elastic hydrogels (covalent crosslinking only) exhibited similar behavior to fibroblasts on viscoelastic hydrogels with equivalent *E* (**Fig. S7**).

**Fig. 3:**
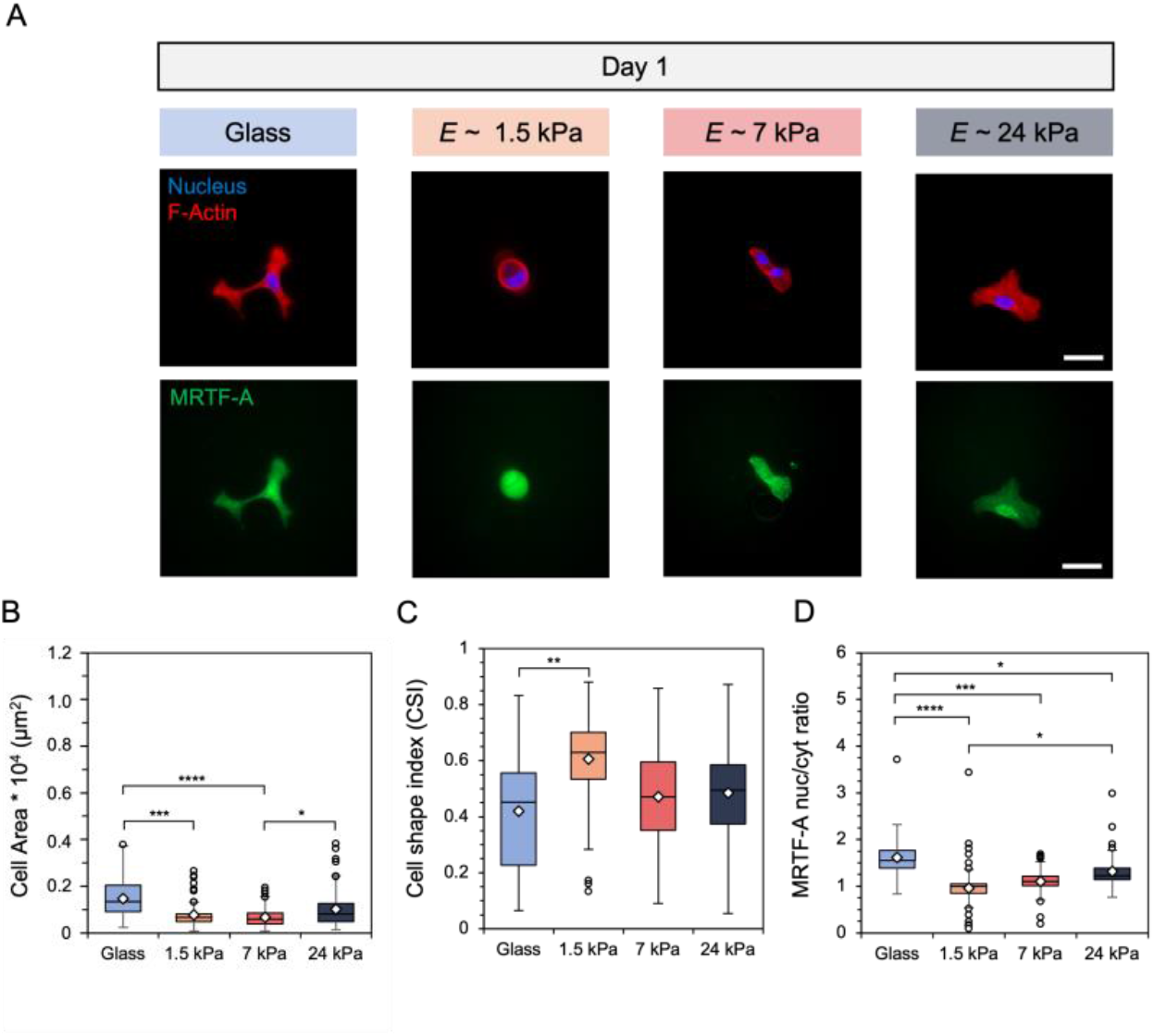
A) Representative images of fibroblasts cultured on glass and hydrogels of E ∼ 1.5, 7, and 24 kPa for 24 hours. Scale bars: 50 μm. Fibroblast B) spread area (μm^2^), C) cell shape index, which measures cell circularity, and D) nuclear localization of myocardin related transcription factor A (MRTF-A) were quantified. *N* = 3 hydrogels per group. ****: P < 0.0001, *** P < 0.0005, ** P < 0.01, * P < 0.05.

### 3.3 Fibroblast global DNA methylation and chromatin condensation levels display time-dependent changes during hydrogel culture

To study how substrate mechanics affected nuclear remodeling events, fibroblasts were cultured atop hydrogels along with glass controls for a period of up to 9 days. Quantification of nuclear metrics included global nuclear DNA methylation intensity, as measured by 5-methylcytosine (5-mC), and chromatin condensation percentage (CCP). After 1 day of culture, fibroblasts displayed significantly greater DNA methylation intensity and significantly lower CCP, as shown by the inset pixelated edges within the nuclear images, on the two stiffer hydrogel groups (**Fig. 4A**). Indeed, quantification showed a two- and five-fold increase in DNA methylation staining intensity for fibroblasts on the 7 and 24 kPa hydrogels, respectively, when compared to the 1.5 kPa hydrogel (**Fig. 4B**). Further, CCP analysis reported a significant decrease in condensed chromatin, from an average of ∼ 17% on the 1.5 kPa hydrogel to 11% and 12% for the 7 and 24 kPa hydrogels, respectively (**Fig. 4C**).

**Fig. 4:**
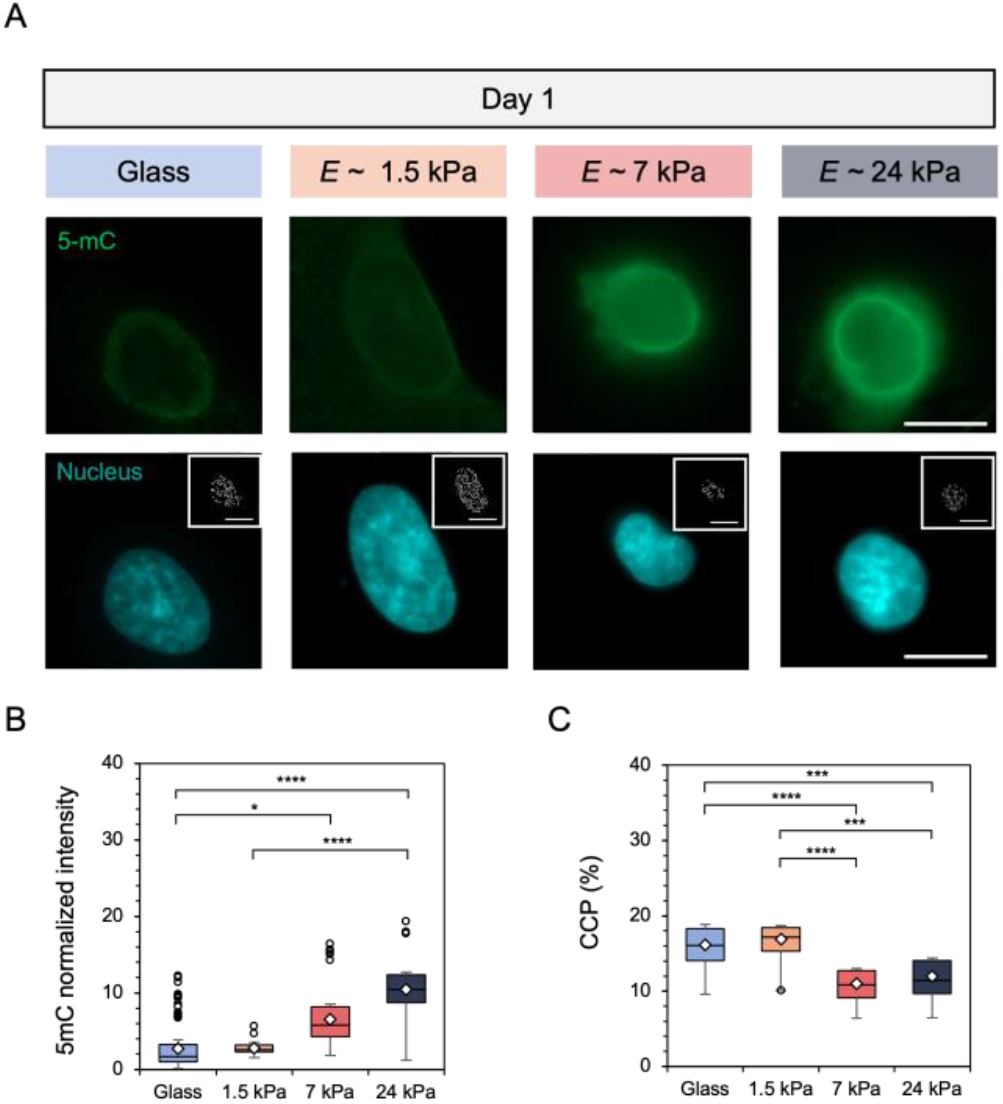
A) Representative images of fibroblast global DNA methylation as indicated by 5-mC (green) staining and nuclei (DAPI) on glass and hydrogels following culture for 24 hours. Inset images represent pixelated edges within the nucle*i*, used to quantify chromatin condensation percentage (CCP). Scale bars: 10 μm. Nuclear metrics were measured by B) global DNA methylation intensity within the nucleus and C) the CCP. *N* = 3 hydrogels per group. **** P < 0.0001, *** P < 0.0005, * P < 0.05.

When fibroblasts underwent longer culture atop the hydrogels, DNA methylation intensity showed a dramatic reduction for all groups by day 9 except the stiffest hydrogel, which showed only a faint nuclear stain, and high levels of chromatin condensation for all groups were observed in the inset images within the nuclear stains (**Fig. S8A**). Quantification of 5-mC nuclear intensity confirmed the decline in DNA methylation over time where, after day 5, all groups report similar levels of methylation (**Fig. S8B)**. Further, chromatin analysis showed increasing condensation over time, with similar levels of condensation for all substrate groups reported after day 5 as well (**Fig. S8C**). To test whether viscoelasticity contributed to the changes observed in nuclear behavior, fibroblasts underwent culture on 1.5 kPa elastic hydrogels compared to 1.5 kPa viscoelastic hydrogels. No distinct differences in DNA methylation or CCP were found (**Fig. S9**).

### 3.4 Hydrogels with engineered in situ secondary crosslinking enable hydrogel stiffening in the presence of cells

After evaluating fibroblast response on mechanically static hydrogels over different culture lengths, we next sought to recapitulate the dynamic progression of fibrosis by assessing fibroblast response on hydrogels amenable to *in situ* stiffening. Due to the efficiency of thiol-ene click reactions, initially compliant viscoelastic hydrogels with *E* ∼ 1.5 kPa can form secondary crosslinks under subsequent exposure to UV light after additional photoinitiator and dithiol crosslinker are added. This allows for hydrogels that can stiffen at user-defined timepoints in the presence of cells to mimic the progressive stiffening nature of fibrotic disease. Rheology time sweeps illustrate the formation of initially soft hydrogels after one dose of UV light, followed by further covalent network crosslinking to stiffen the hydrogel to *E* ∼ 24 kPa after a second UV light exposure (**Fig. 5A**). Notably, the exposure to light and/or photoinitiator does not affect cell spreading behavior and is not cytotoxic (**Fig S10**). This hydrogel system was then used to probe fibroblast response following hydrogel stiffening at 1 or 7 days post-seeding onto the initially compliant *E* ∼ 1.5 kPa hydrogel (**Fig. 5B**).

**Fig. 5:**
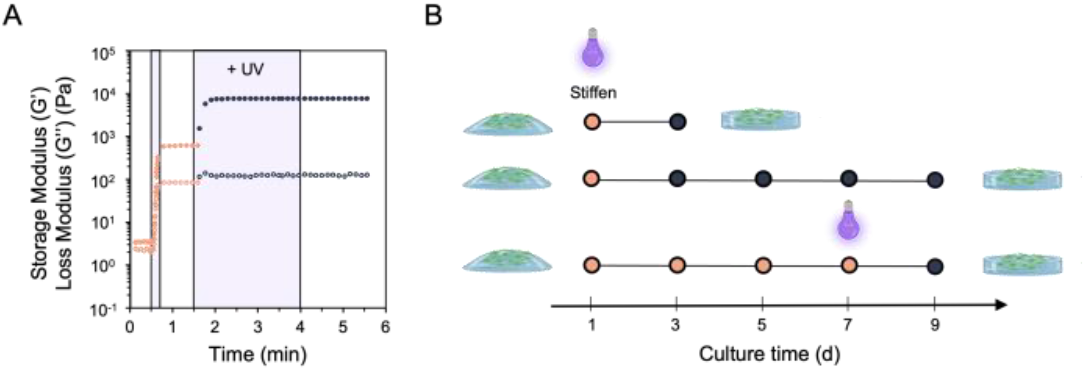
A) Rheology of initially soft, viscoelastic hydrogels that can undergo secondary crosslinking following subsequent doses of light exposure to match the mechanics of the stiffest hydrogel group. B) Using this hydrogel system, fibroblasts were cultured atop the soft, viscoelastic hydrogel for a period of either 1 or 7 days prior to experiencing *in situ* secondary stiffening, then fixed and stained to quantify cell shape and nuclear metrics.

### 3.5 Human lung fibroblast phenotypic and nuclear behavior rapidly respond to early, but not late, hydrogel stiffening

Fibroblasts underwent culture on the initial *E* ∼ 1.5 kPa hydrogel for either 1 or 7 days prior to *in situ* stiffening. Cell response was compared to mechanically static hydrogels of *E* ∼ 1.5 kPa or 24 kPa to match pre- and post-stiffening mechanics respectively. Stiffening the hydrogel after 1 day resulted in increased fibroblast spreading, F-actin stress fiber formation, and more intense nuclear MRTF-A staining after 2 days (**Fig. 6A**). Quantification showed an increase in the range of cell spread area, with some areas reaching above 0.4 × 10^4^ μm^2^ (**Fig. 6B**). Fibroblasts also became ∼ 30% more elongated than the mechanically static 1.5 kPa control, as measured by averages of the cell shape index (**Fig. 6C**). While no statistically significant difference was observed for changes in MRTF-A nuclear localization for fibroblasts that experienced hydrogel stiffening, quantification showed an upward trend toward the mechanically static 24 kPa hydrogel group (**Fig. 6D**). Similar phenotypic results were obtained for fibroblasts that experienced stiffening after 1 day but were cultured for a total of 9 days (**Fig. S11**).

**Fig. 6:**
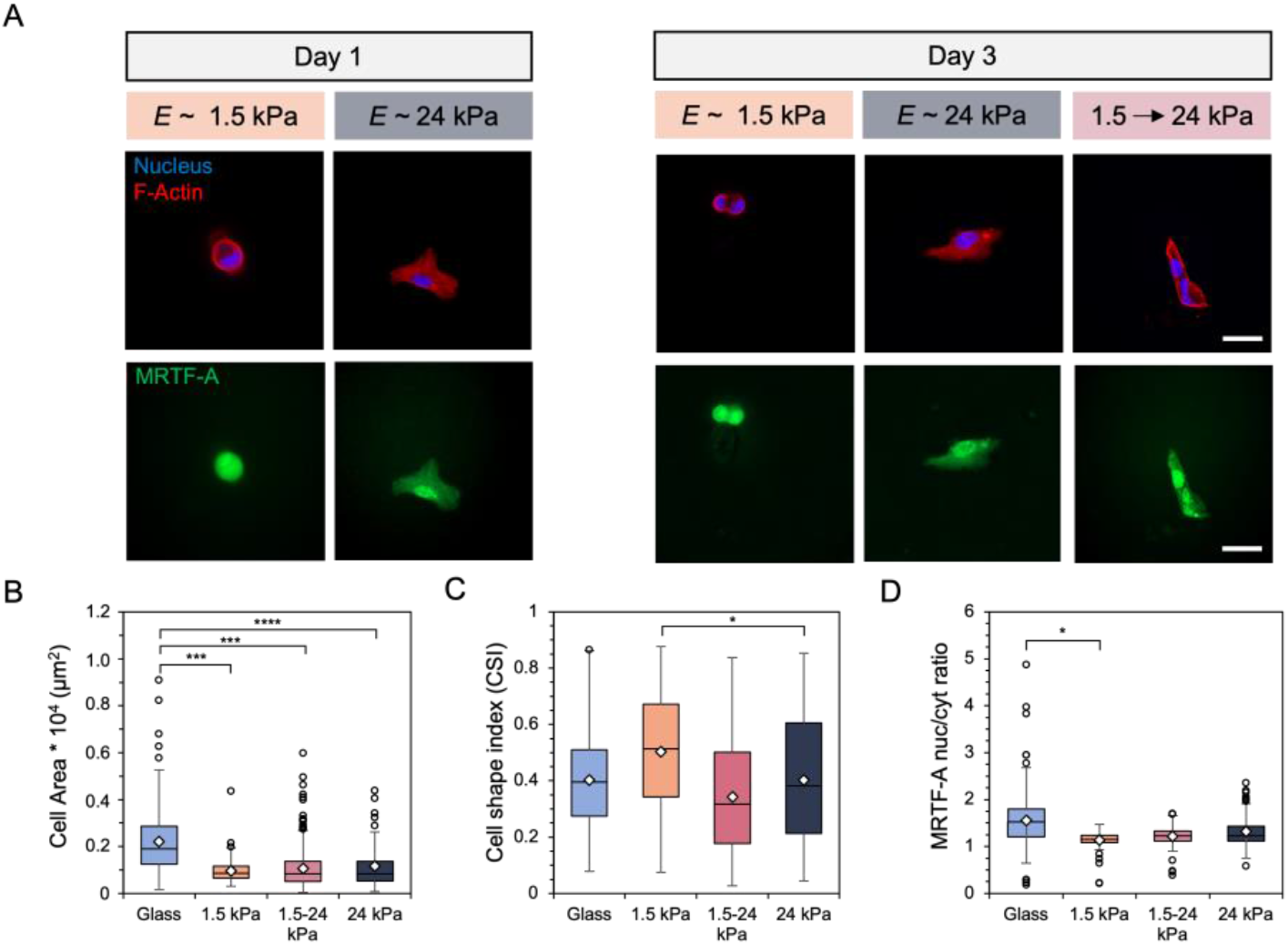
A) Representative images of fibroblasts after (left) one day of culture on 1.5 or 24 kPa mechanically static hydrogels and (right) after 3 days on either the mechanically static hydrogels or a hydrogel that was stiffened after 1 day. Scale bars: 50 μm. *F*ibroblast B) spread area (μm^2^), C) cell shape index, and D) MRTF-A nuclear localization were quantified. *N* = 3 hydrogels per group. **** P < 0.0001, *** P < 0.0005, * P < 0.05.

Similarly, fibroblast nuclear behavior appeared to respond to changes in substrate mechanics following early stiffening events. DNA methylation staining increased in intensity while the amount of chromatin shown within the nuclear inset images appeared to decrease after 2 days on the stiffened hydrogel (**Fig. 7A**). Analysis of nuclear 5-mC intensity showed an increase in global DNA methylation compared to the static 1.5 kPa hydrogel following a 3 day total culture, although no statistically significant differences were reported (**Fig. 7B**). Similarly, CCP results indicated a decrease in chromatin condensation toward results obtained for fibroblasts cultured on mechanically static 24 kPa hydrogels for 3 days (**Fig. 7C**). Longer culture times on the stiffened hydrogel such that fibroblasts experienced a 9 day total culture led to nuclear behaviors resembling results obtained for all mechanically static groups after 9 days, with reduced global DNA methylation and high percentages of condensed chromatin (**Fig. S12**).

**Fig. 7:**
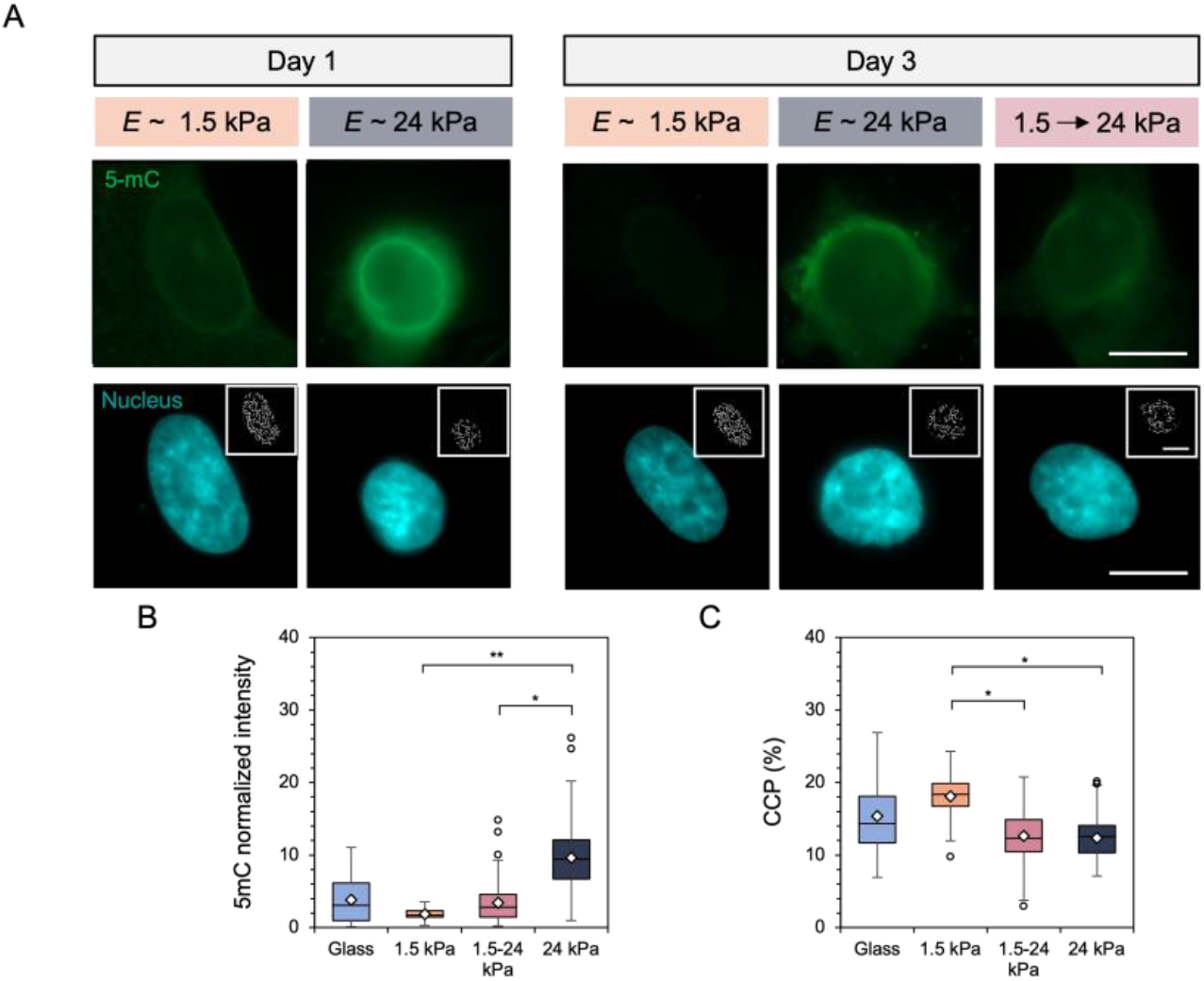
A) Representative images of fibroblast global DNA methylation as indicated by 5-mC (green) staining and nuclei (DAPI) after (left) one day of culture on 1.5 or 24 kPa mechanically static hydrogels and (right) after 3 days on either the mechanically static hydrogels or a hydrogel that was stiffened after 1 day. Inset images represent pixelated edges within the nuclei, used to quantify chromatin condensation percentage (CCP). Scale bars: 10 μm. Nuclear metrics were measured by B) global DNA methylation intensity within the nucleus and C) the CCP. *N* = 3 hydrogels per group. ** P < 0.01, * P < 0.05.

Unlike fibroblast response to early (1 day) stiffening, when fibroblasts experienced late *in situ* stiffening after 7 days there were no significant changes in DNA methylation or chromatin condensation observed (**Fig. S13A**). Analysis of both 5-mC nuclear intensity and chromatin condensation showed similar levels to the mechanically static hydrogel groups as well, suggesting a time-dependent effect on the responsiveness of fibroblast nuclear remodeling following dynamic mechanical perturbations (**Fig. S13B, C**). Furthermore, fibroblast phenotype appeared to trend toward the behaviors observed and measured for static 24 kPa hydrogels (**Fig. S14**).

## 4. Discussion

Hyaluronic acid (HA) was chosen as the hydrogel backbone since it is amenable to chemical modification with a variety of functional groups to enable diverse crosslinking mechanisms^50^. This allows fabrication of HA hydrogels with tailored mechanical properties, like stiffness and viscoelasticity, similar to both normal and fibrotic tissues, as reported by our group and others^16, 27, 28, 50^. Norbornene modification to HA (NorHA) afforded the use of light-mediated thiol-ene click chemistry. Compared to other alkene functional groups that form kinetic chains, like (meth)acrylates, norbornenes have low reactivity with themselves, making them a promising functional moiety for stable and controllable covalent network formation^46^. Modification of HA with β-cyclodextrins enabled crosslinking through physical interactions with adamantane (Ad) peptides. The Ad peptides included thiol-containing cysteines which could crosslink with available norbornenes, so that Ad could be tethered to the polymer backbone. Supramolecular guest-host crosslinks could then form between the Ad (guest), which has a high affinity toward the hydrophobic cavity of β-cyclodextrin (host), groups (Fig. 1). This system is highly tunable where, once the initial stiffness and viscoelastic properties are determined, introduction of further covalent crosslinks through thiol-norbornene click chemistry increases the storage modulus (G’) while holding the loss modulus (G’’) relatively constant (Fig. 2). Notably, the softest hydrogel formulation used in this work showed a G” within an order of magnitude of G’, which is a property of native viscoelastic lung tissue^26, 51^. This system enabled the formation of hydrogels with a wide range of stiffnesses, from G’ ∼ 0.5 kPa to G’ ∼ 8 kPa, by simply adjusting the dithiol concentration or UV exposure time (Fig. 2A). The increase in storage modulus also widened the gap between the storage and loss moduli, forming physiologically relevant hydrogels that were stiffer and more elastic in a highly tunable manner (Fig. 2B)^51^. Further characterization through frequency sweeps and stress relaxation testing showed that the softest hydrogel displayed a greater increase in loss moduli with increased frequency and stress relaxation following an applied strain. This was likely due to the higher relative content of physical guest-host interactions in the network compared to the two stiffer hydrogel formulations (Fig. 2C,D).

After mechanical characterization confirmed the ability to form hydrogels with independently tunable stiffness and viscoelasticity, we studied how human lung fibroblasts responded to the three hydrogel groups, as well as a non-hydrogel glass control, for a culture period of 9 days. After just one day, fibroblasts began displaying distinct morphologies in response to these substrates, with greater actin stress fiber organization, spreading, and elongation with respect to substrate stiffness (Fig. 2A top, B, C). Furthermore, the translocation of myocardin related transcription factor A (MRTF-A) into the nucleus also increased with increasing substrate stiffness. Following cell-matrix contact, MRTF-A, which is a transcriptional coactivator, moves into the nucleus and interacts with the serum response factor (SRF). SRF is a transcription factor that plays a key role in the upregulation of *Acta2*, which is one of the hallmark genes involved in fibroblast activation from a quiescent state^52-55^. All metrics of cell shape and MRTF-A nuclear-to-cytosol ratios remained constant throughout the remainder of the culture (Fig. S5) and do not show dependence on substrate viscoelasticity at lower stiffnesses (Fig. S6). These results indicate that cell phenotype undergoes changes within hours of experiencing different substrate mechanics and that these behaviors remain largely unchanged over the course of 9 days. The data reported here also corroborate other findings in cellular phenotype with respect to substrate stiffness^6, 7, 56^.

Compared to our collective understanding of phenotypic responses^3, 15, 16, 28, 57, 58^ and even MRTF-A^7, 27, 55, 59^ localization with respect to substrate mechanics and culture time, there are very few reports of cell nuclear metrics^37, 38, 40-42^, and even less so for global DNA methylation^60^ analyses in response to substrate mechanics over time. We next investigated fibroblast nuclear responses to the glass and three hydrogel substrates for the same culture duration. Similar to the fibroblast shape and MRTF-A metrics evaluated, the nuclear markers of global DNA methylation, as measured by 5-methylcytosine (5-mC) intensity, along with the amount of condensed chromatin correlated with substrate mechanics after 1 day of culture. DNA methylation showed greater intensity with increasing hydrogel stiffness, while the percentage of condensed chromatin (CCP) was reduced, or more open and available for downstream transcriptional processes (Fig 4). DNA methylation has been implicated in driving cell differentiation and others have also reported increased methylation with respect to substrate stiffness, supporting our findings^45^. Similarly, other groups also reported reduced CCP values for cells cultured on stiffer matrices, implicating this more open structure in greater transcriptional activity, which also corroborates our findings^37, 38, 40^. Notably, fibroblast DNA methylation and CCP do not respond to glass culture in the same way that they do on stiff hydrogels, which is believed to be due to their initial culture and expansion on tissue culture plastic of similar stiffness (GPa) prior to these *in vitro* studies^37^.

Interestingly, both nuclear markers were found to display culture time-dependent trends such that over the course of 9 days, fibroblasts cultured on the two stiffer hydrogel groups showed reduced DNA methylation and increased CCP to basal levels seen in the glass and softest hydrogel group (Fig. 5). Furthermore, these nuclear metrics do not seem to depend on substrate viscoelasticity, suggesting that stiffness plays a greater role in global DNA methylation and chromatin condensation states (Fig. S7). Others have also noted similar time-dependent changes in CCP, suggesting that after a certain time cells obtain a ‘persistent’ state where, once CCP levels match that of the softest mechanics, cells become less sensitive to mechanical changes^40^. However, to the best of our knowledge, this is the first time that DNA methylation has been shown to display a time-dependent response to substrate stiffness.

To further explore how culture timing affects fibroblast response to mechanical cues, we fabricated hydrogels that underwent *in situ* stiffening at user-defined timepoints by exploiting the ability to introduce secondary thiol-ene crosslinks in our hydrogel system. By swelling in additional dithiol crosslinker and photoinitiator following initial network formation of *E* ∼ 1.5 kPa hydrogels, a second UV light exposure enabled further covalent crosslinking to produce *E* ∼ 24 kPa hydrogels. This enabled us to initially culture fibroblasts on the *E* ∼ 1.5 kPa stiffness, then expose them to stiffer mechanics at early or late timepoints (Fig. 6). Day 1 and day 7 were chosen as the early and late stiffening timepoints, respectively, as fibroblast nuclear response seemed to begin changing after ∼ 5 days of culture (Fig. 5B, C). As expected, early stiffening at day 1 resulted in immediate changes to both fibroblast spreading as well as nuclear reorganization toward trends observed for the mechanically static *E* ∼ 24 kPa control (Fig. 7, 8). However, after 7 days of culture on *E ∼* 1.5 kPa hydrogels prior to stiffening, fibroblasts maintained low levels of DNA methylation and highly condensed chromatin, even though the substrate mechanics had been increased (Fig. 9). These results highlight that fibroblast activation, which can be observed within 1 day through phenotypic analysis, can take ∼ 5 days to become persistent and is implicated by changes in DNA methylation and chromatin condensation.

Overall, this work examined how human lung fibroblast spreading and nuclear reorganization respond to both static and dynamic mechanical signals over time using a hydrogel platform mimicking the stiffness and viscoelastic properties of healthy and increasingly fibrotic lung tissue. Our results show that fibroblast spreading and MRTF-A nuclear localization rapidly respond to changes in stiffness and remain stable over time, but nuclear metrics (DNA methylation, chromatin condensation) are more dependent on the length of time spent on the substrate. This could have implications on future studies aiming to better understand, or even reverse, fibroblast activation. Future experiments may look to inhibit specific types of DNA methylation to investigate its role in chromatin condensation and fibroblast de-activation as a potential target for reversing fibrosis, along with considering the response of other cell types/sources, like primary cells or fibroblasts derived from fibrotic tissue, to better understand cell-specific behaviors.

## Supporting Information

^1^H NMR spectra for the hydrogel components, MALDI for the adamantane peptide, and additional data from the cell culture experiments can be found in the Supporting Information.

## Acknowledgments

This work was supported by the NSF (CAREER DMR/BMAT 2046592) and NIH (R35GM138187). The content is solely the responsibility of the authors and does not necessarily represent the official views of the National Institutes of Health. The authors would also like to acknowledge Drew Miller for helpful discussions regarding epigenetic investigations.

