## Supplemental Information for "Hydrogel mechanics regulate fibroblast DNA methylation and chromatin condensation"

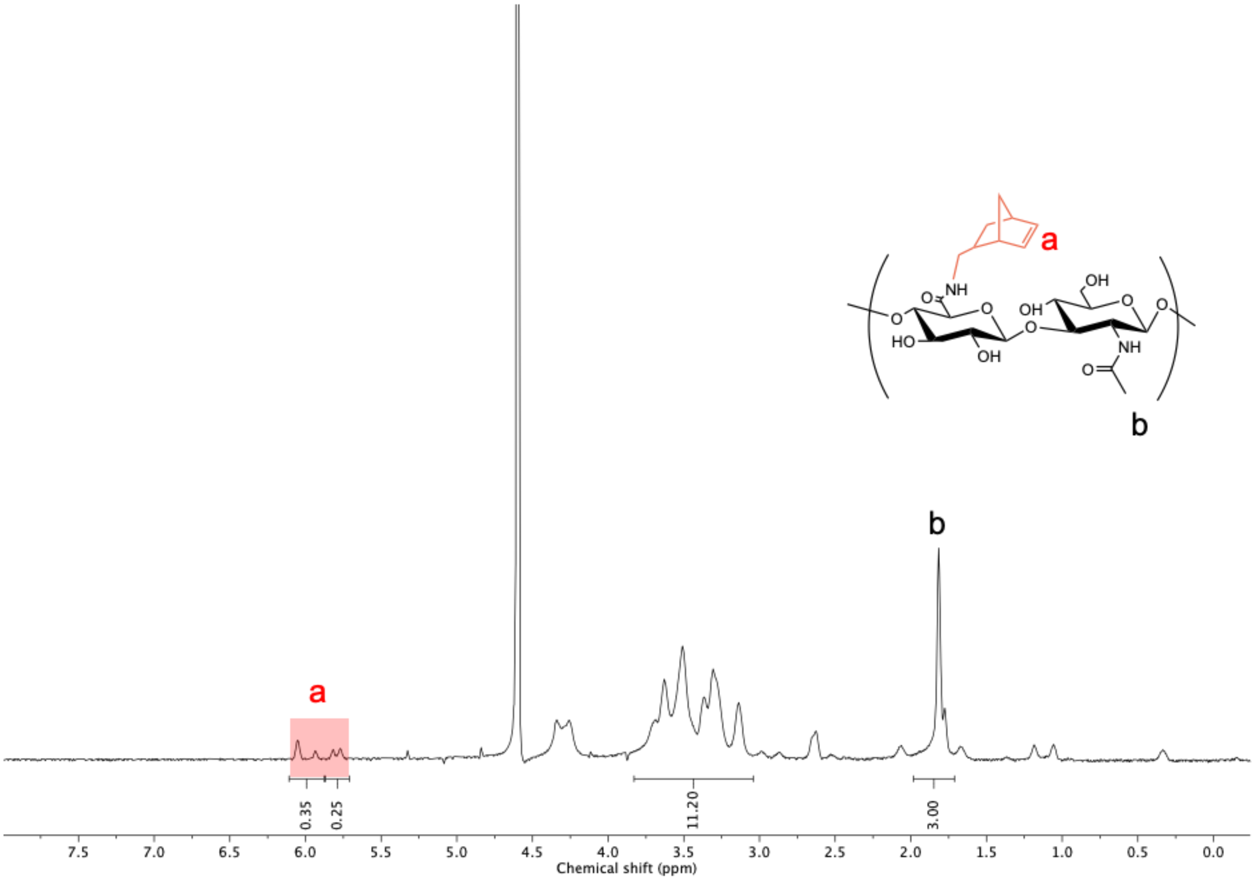


**Figure S1**. $^1$H NMR spectrum of norbornene-modified hyaluronic acid (NorHA). The degree of modification was determined to be 30% as indicated by integration of the peaks at δ=5.75, 6.05, and 6.17 (2H, shaded red) relative to the N-acetyl on HA (3H).

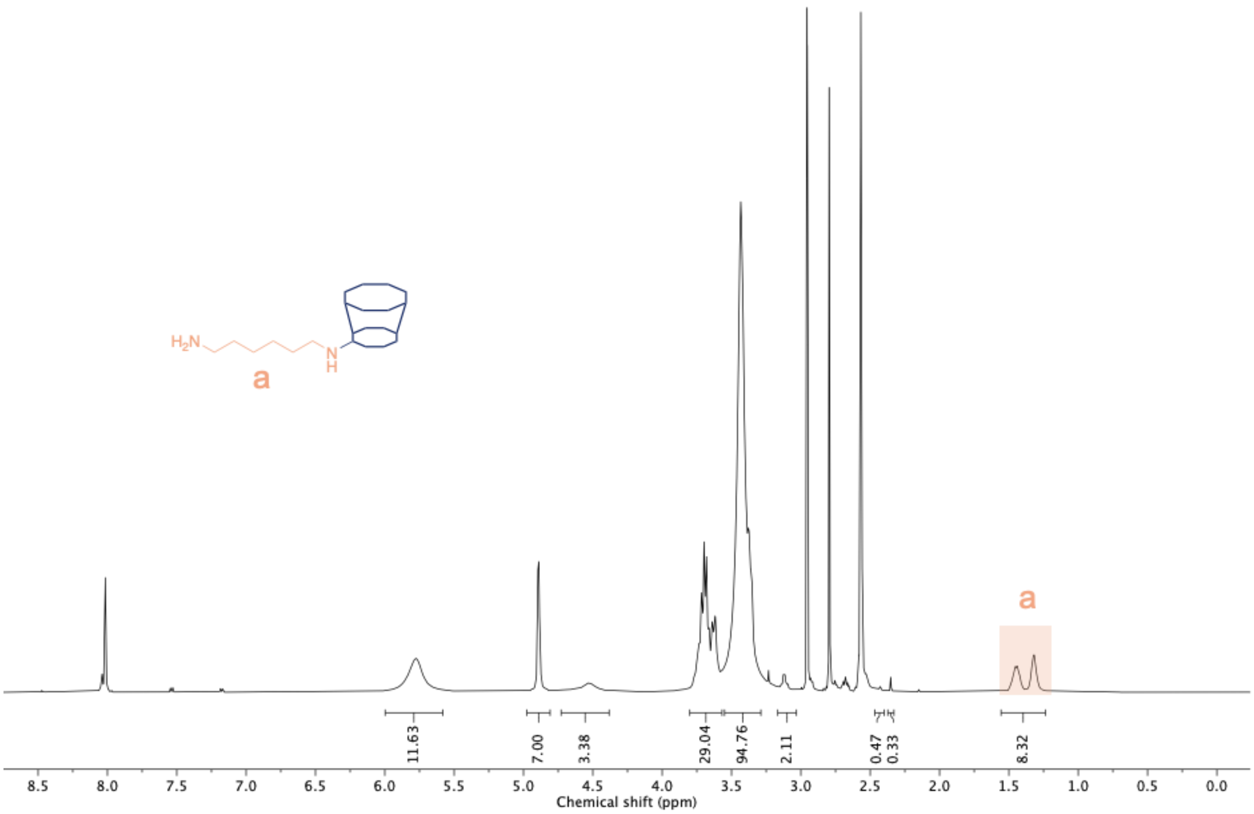


**Figure S2**. [1]H NMR spectra of β-cyclodextrin hexamethylene diamine (β-CD-HDA). Modification of HDA was determined to be 69% as determined by the peaks at δ=1.14-1.6 ppm (12H).

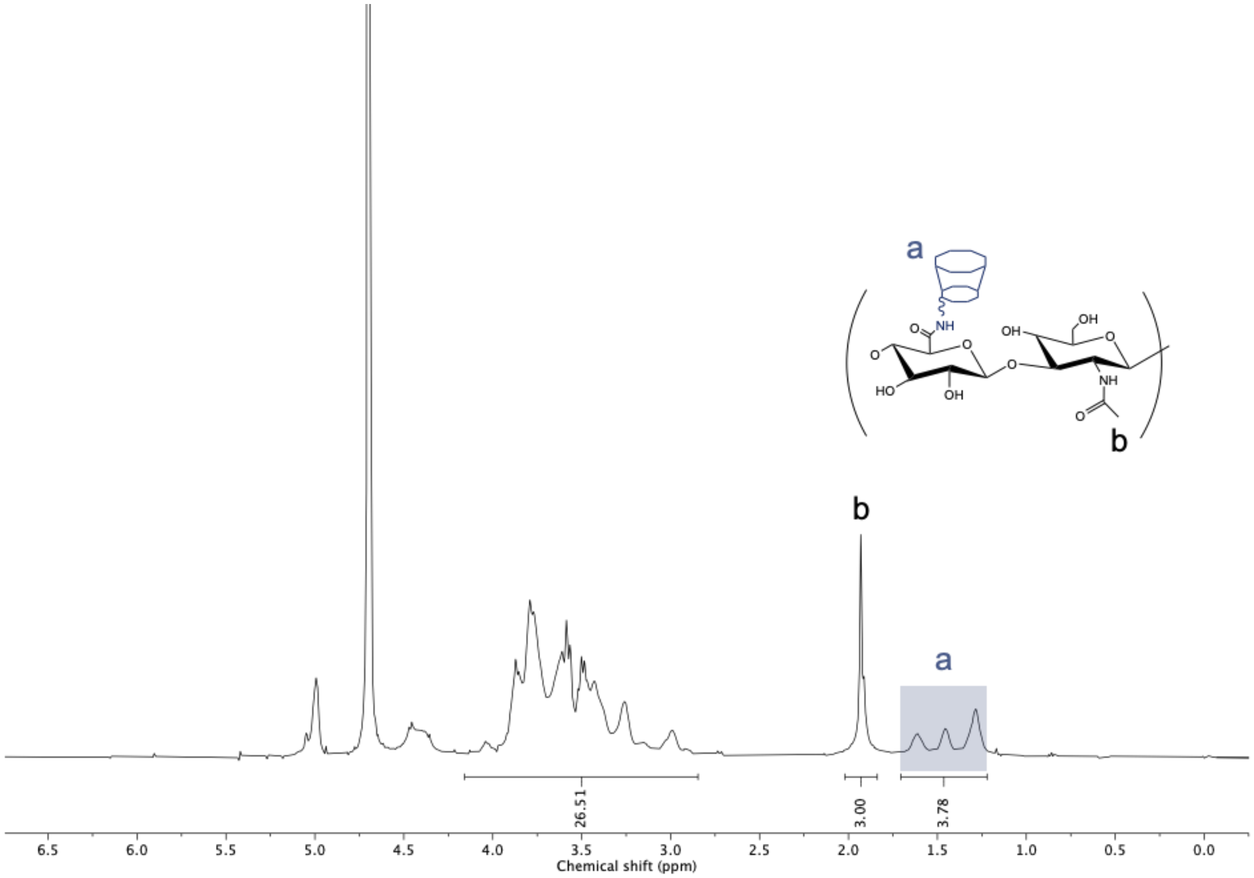


**Figure S3**. $^1$H NMR spectra of β-cyclodextrin-modified hyaluronic acid (CD-HA). The degree of modification was found to be 31% as determined by integration of the peaks at δ=1.23-1.68 ppm (12H) relative to the N-acetyl on HA (3H).

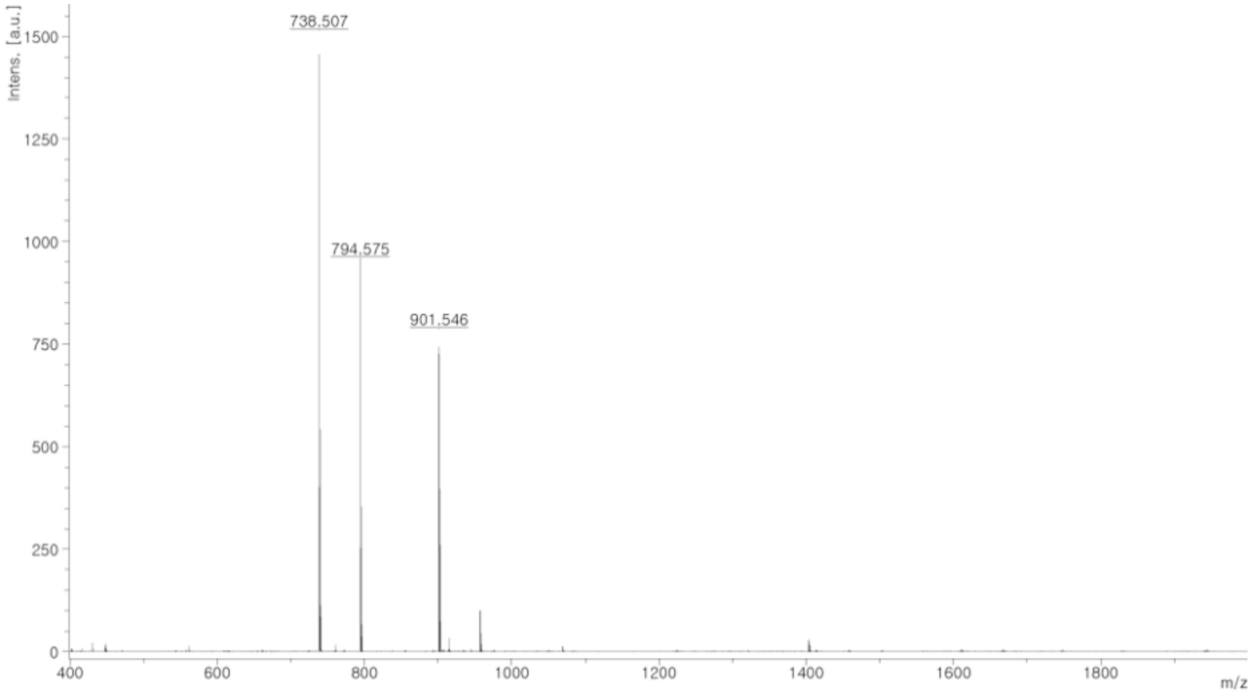

**Figure S4**. MALDI spectrum of 1-adamantaneacetic acid-KKKCG (adamantane peptide). Expected mass: 738.6 g/mol. Actual mass: 738.5 g/mol.

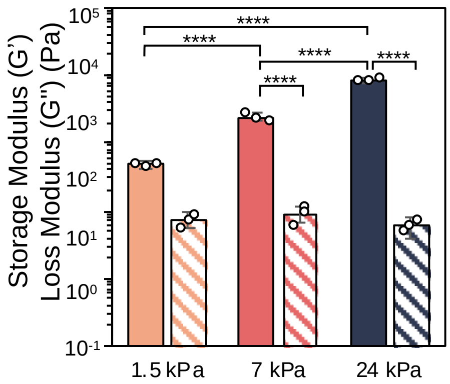


**Figure S5**. Quantification of the storage (solid filled) and loss (patterned) modulus for each hydrogel stiffness. Data reported as mean ± s.d. $N$ = 3 hydrogels per group. **** $P < 0.0001$.

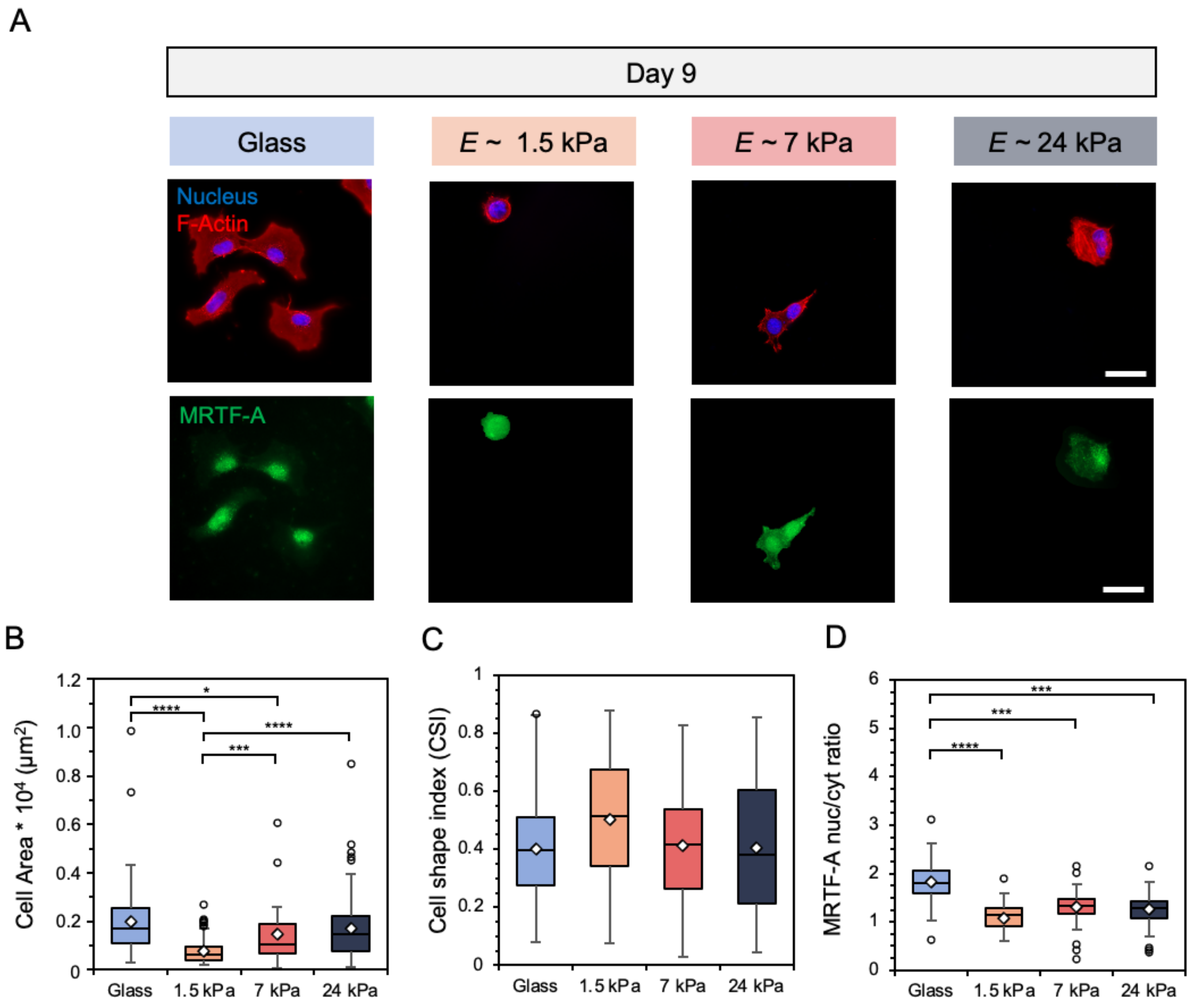


**Figure S6**. **A**) Representative images of fibroblasts cultured on glass and hydrogels of E ~ 1.5, 7, and 24 kPa for 9 days. Scale bars: 50 µm. Fibroblast **B**) spread area (µm$^2$), **C**) cell shape index, which measures cell circularity, and **D**) nuclear localization of myocardin related transcription factor A (MRTF-A) were quantified. $N$ = 3 hydrogels per group. ****: P < 0.0001, *** P < 0.0005, ** P < 0.01, * P < 0.05.

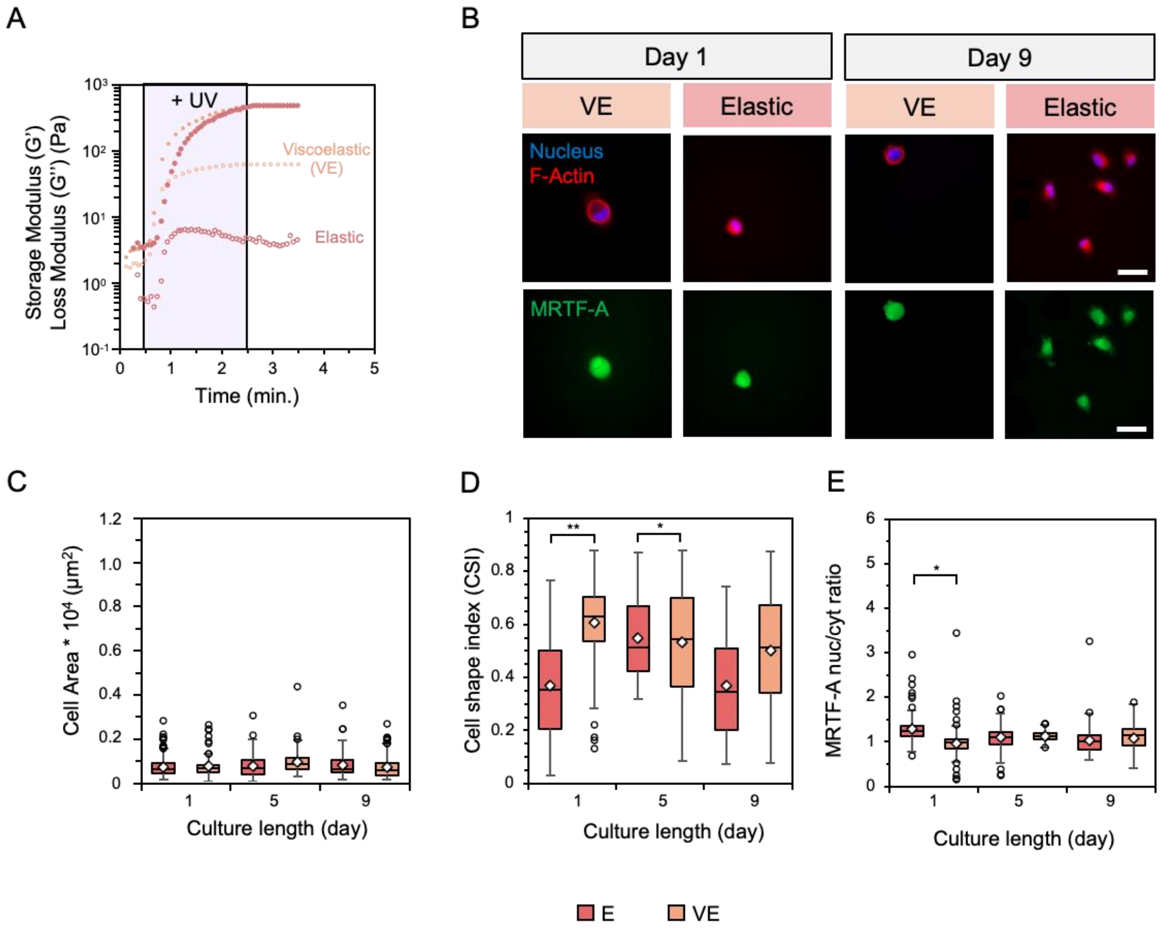


**Figure S7**. **A**) *In situ* rheology of soft viscoelastic (VE) and soft elastic (E) hydrogels of the same storage modulus (closed circles) but different loss moduli (open circles). **B**) Representative images of fibroblasts cultured on hydrogels following 1 (left) and 9 (right) days of culture. Scale bars: 50 μm. Fibroblast **C**) spread area (μm$^2$), **D**) cell shape index, and **E**) MRTF-A nuclear localization were quantified. $N$ = 3 hydrogels per group. ** $P < 0.01$, * $P < 0.05$.

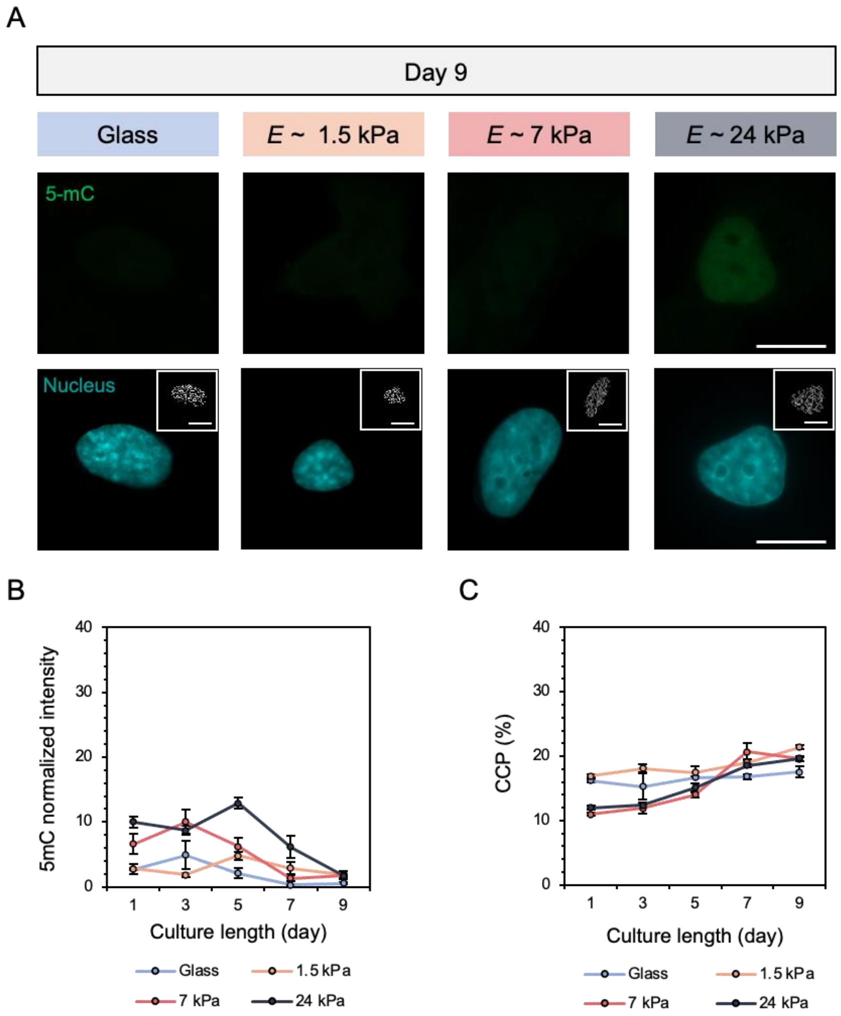


**Figure S8**. **A**) Representative images of fibroblast global DNA methylation as indicated by 5-mC (green) staining and nuclei (DAPI) on glass and hydrogels following culture for 9 days. Inset images represent pixelated edges within the nuclei, used to quantify chromatin condensation percentage (CCP). Scale bars: 10 μm. Nuclear metrics of **B**) normalized global DNA methylation intensity and **C**) CCP were quantified as a function of culture length for a total of 9 days. $N = 3$ hydrogels per group. Data reported as total mean ± s.e.m.

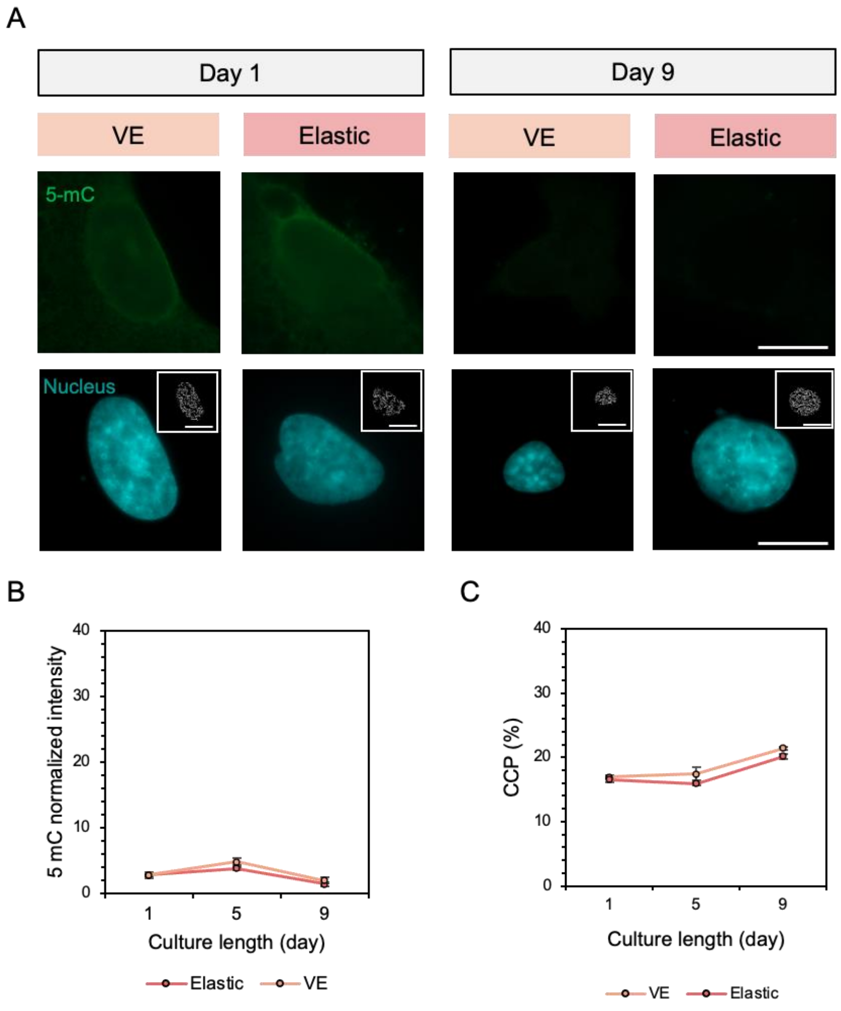


**Figure S9**. **A**) Representative images of fibroblast global DNA methylation as indicated by 5-mC (green) staining and nuclei (DAPI) on viscoelastic and elastic hydrogels with equivalent $E$ ~ 1.5 kPa following 1 (left) and 9 (right) days of culture. Inset images represent pixelated edges within the nuclei, used to quantify chromatin condensation percentage (CCP). Scale bars: 10 µm. Nuclear metrics of **B**) normalized global DNA methylation intensity and **C**) CCP were quantified as a function of culture length for a total of 9 days. $N$ = 3 hydrogels per group. Data reported as total mean ± s.e.m.

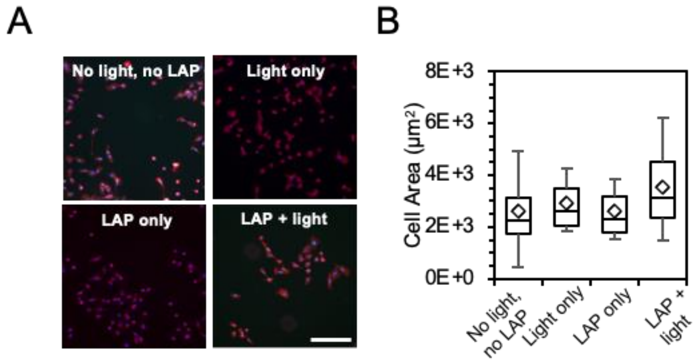


**Figure S10**. **A**) Fibroblast response to UV (365 nm) light, 1 mM LAP, and LAP + UV light, as well as a no light, no LAP control. Scale bar: 200 μm. **B**) Cell spread area quantification 24 h after light and/or LAP exposure showed no statistically significant differences between experimental groups.

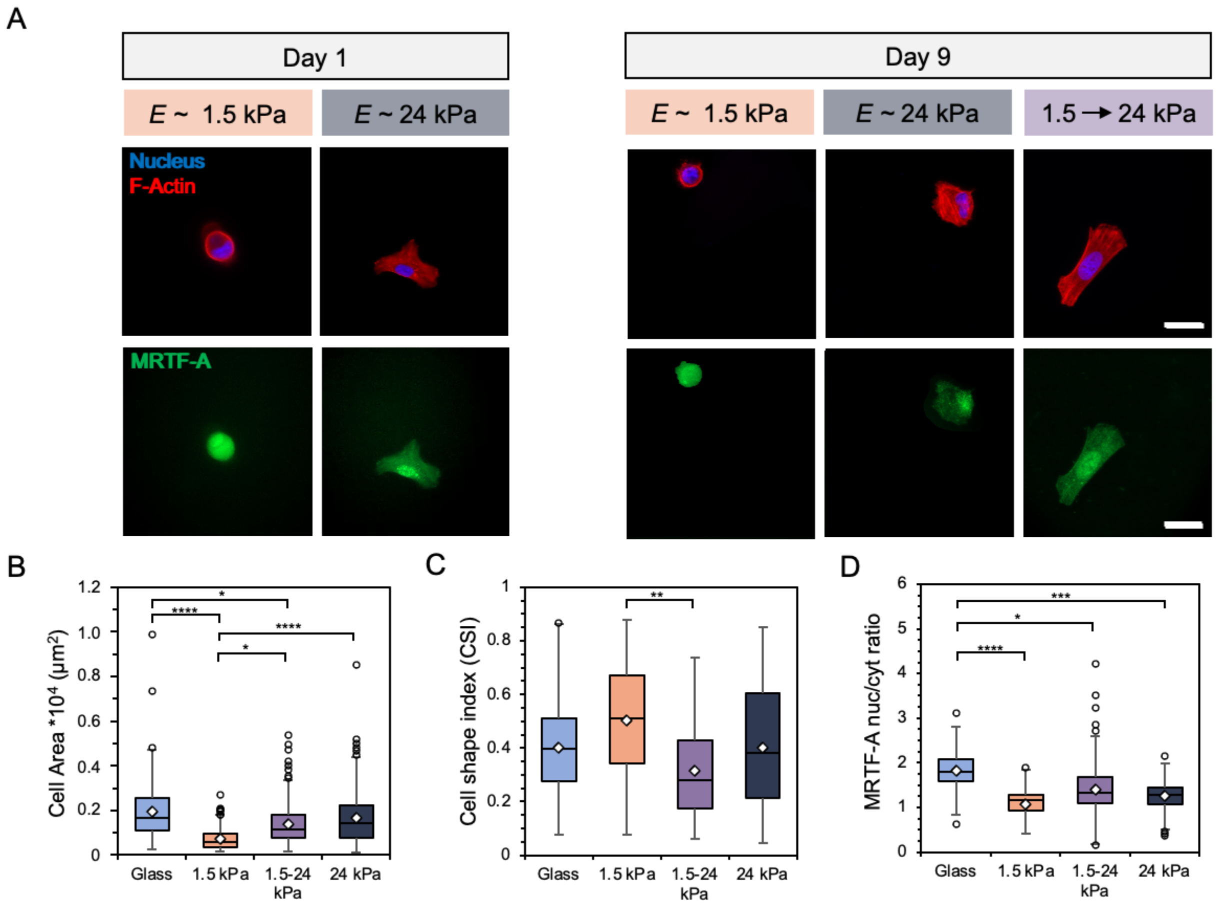


**Figure S11**. **A**) Representative images of fibroblasts after (left) 1 day of culture on E ~ 1.5 or 24 kPa mechanically static hydrogels and (right) after 9 days on either the mechanically static hydrogels or a hydrogel that was stiffened after 1 day. Scale bars: 50 μm. Fibroblast **B**) spread area ($\mu m^2$), **C**) cell shape index, and **D**) MRTF-A nuclear localization were quantified. *N* = 3 hydrogels per group. **** P < 0.0001, *** P < 0.0005, ** P < 0.01, * P < 0.05.

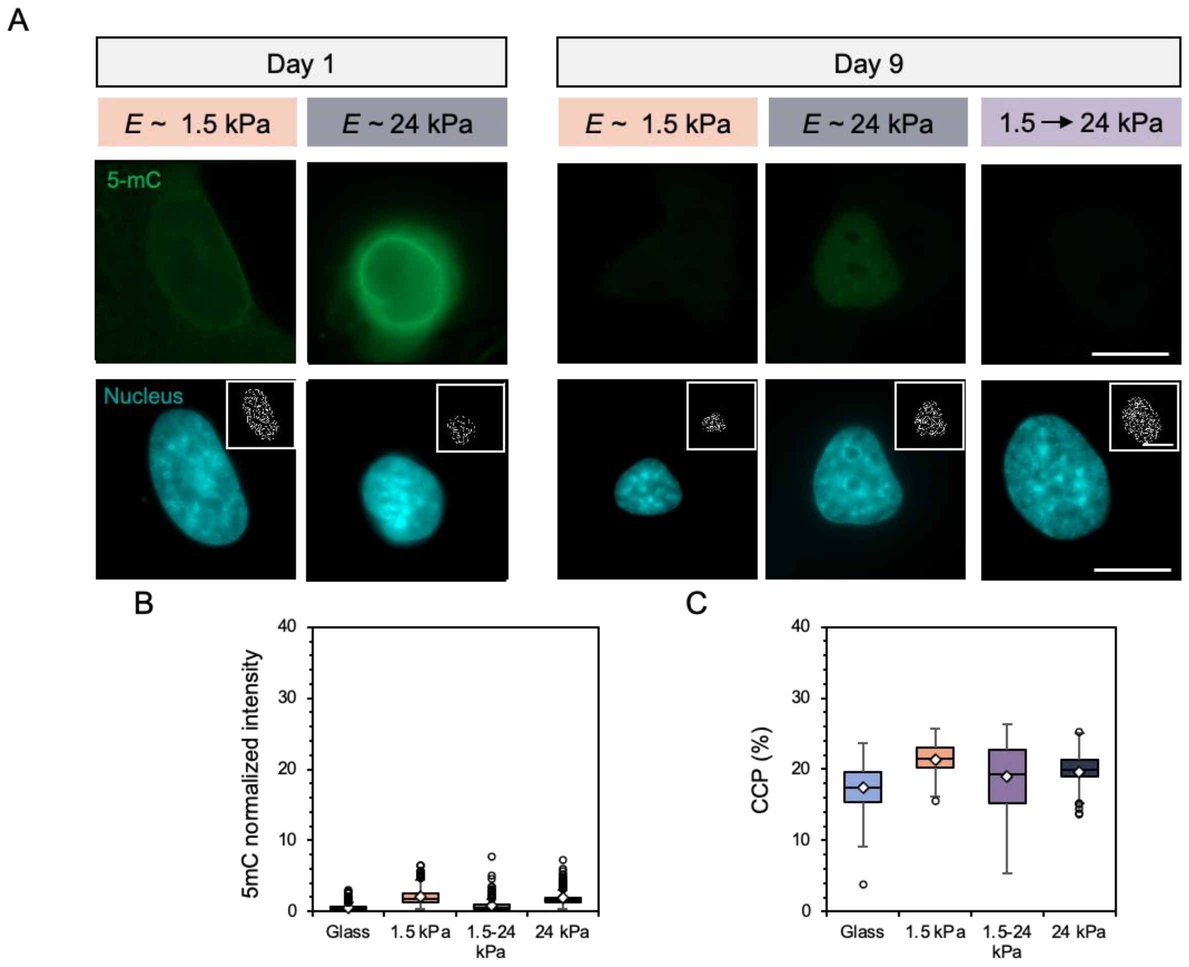


**Figure S12**. **A**) Representative images of fibroblast global DNA methylation as indicated by 5-mC (green) staining and nuclei (DAPI) after (left) one day of culture on E ~ 1.5 or 24 kPa mechanically static hydrogels and (right) after 9 days on either the mechanically static hydrogels or a hydrogel that was stiffened after 1 day. Inset images represent pixelated edges within the nuclei, used to quantify chromatin condensation percentage (CCP). Scale bars: 10 µm. Nuclear metrics were measured by **B**) global DNA methylation intensity within the nucleus and **C**) the CCP. $N$ = 3 hydrogels per group. No statistically significant differences were observed.

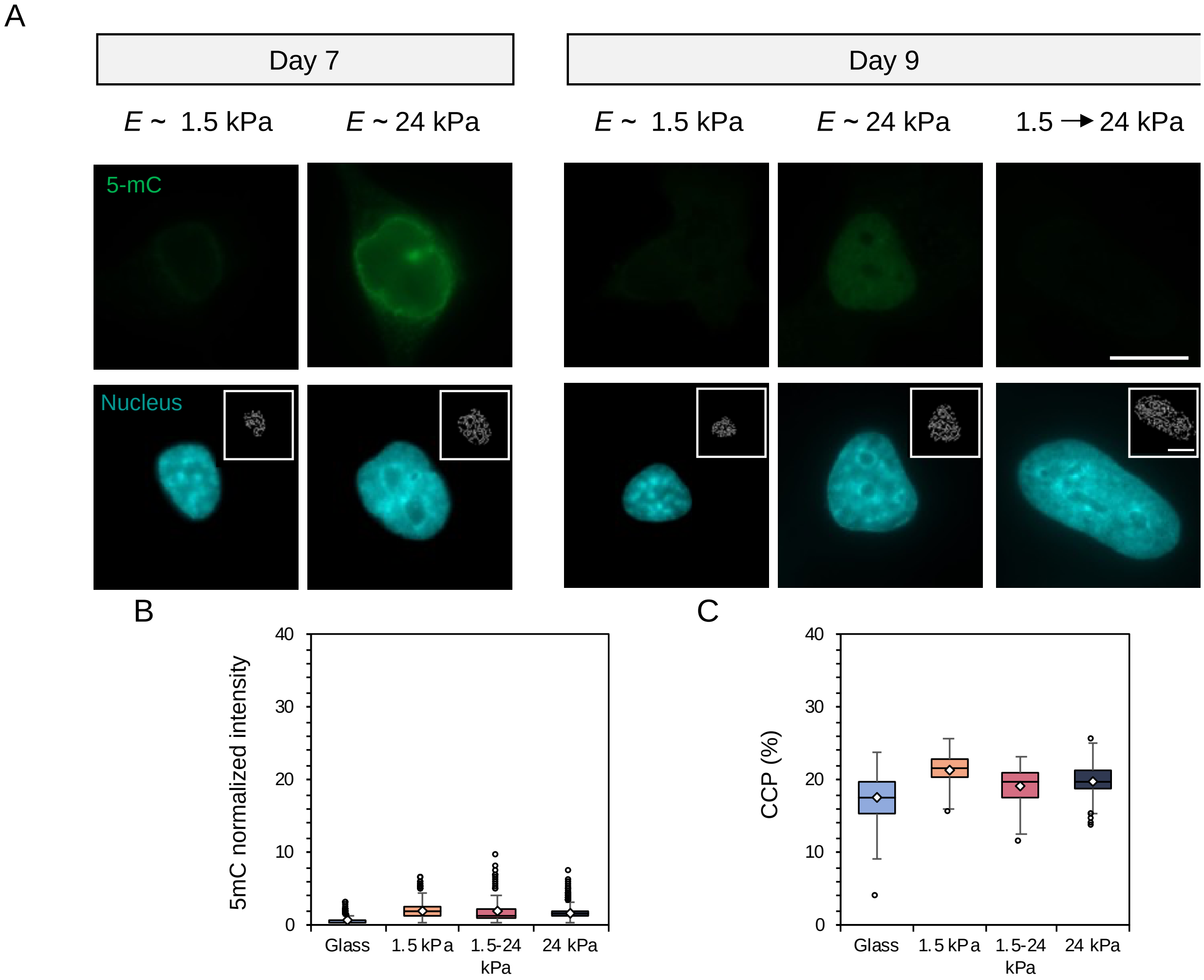


**Figure S13**. **A**) Representative images of fibroblast global DNA methylation as indicated by 5-mC (green) staining and nuclei (DAPI) after (left) one day of culture on E ~ 1.5 or 24 kPa mechanically static hydrogels and (right) after 3 days on either the mechanically static hydrogels or a hydrogel that was stiffened after 1 day. Inset images represent pixelated edges within the nuclei, used to quantify chromatin condensation percentage (CCP). Scale bars: 10 µm. Nuclear metrics were measured by **B**) global DNA methylation intensity within the nucleus and **C**) the CCP. $N = 3$ hydrogels per group. No statistically significant differences were observed.

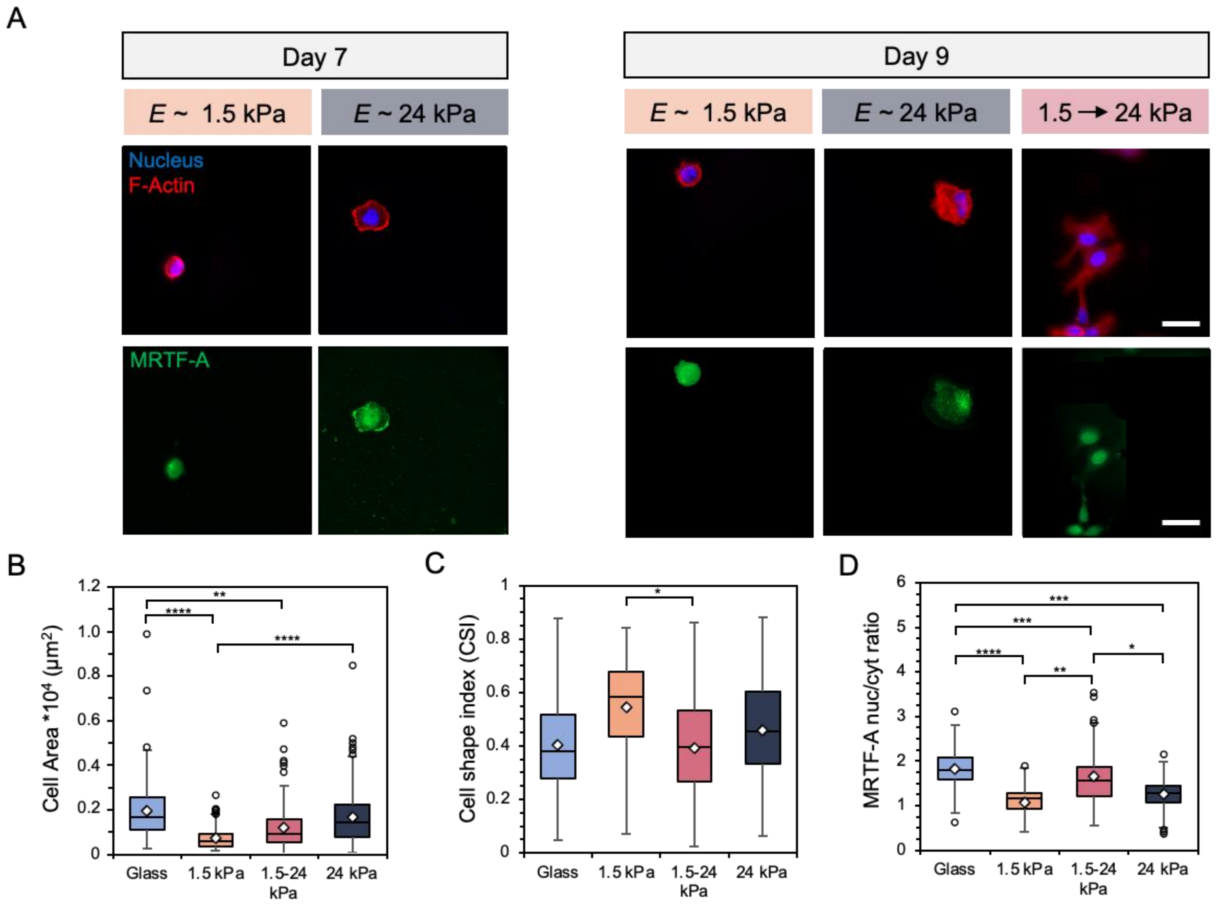


**Figure S14**. **A**) Representative images of fibroblasts after (left) 7 days of culture on E ~ 1.5 or 24 kPa mechanically static hydrogels and (right) after 9 days on either the mechanically static hydrogels or a hydrogel that was stiffened after 7 days. Scale bars: 50 µm. Fibroblast **B**) spread area ($\mu m^2$), **C**) cell shape index, and **D**) MRTF-A nuclear localization were quantified. $N$ = 3 hydrogels per group. **** $P < 0.0001$, *** $P < 0.0005$, ** $P < 0.01$, * $P < 0.05$.
